# Simulating Freely-diffusing Single-molecule FRET Data with Consideration of Protein Conformational Dynamics

**DOI:** 10.1101/2021.01.19.427359

**Authors:** James Losey, Michael Jauch, Axel Cortes-Cubero, Haoxuan Wu, Adithya Polasa, Stephanie Sauve, Roberto Rivera, David S. Matteson, Mahmoud Moradi

**Affiliations:** Department of Chemistry and Biochemistry, University of Arkansas, Fayetteville, AR 72701, U.S.A.; Department of Statistics, Florida State University, Tallahassee, FL 32306, U.S.A.; Department of Mathematical Sciences, University of Puerto Rico, Mayaguez, Puerto Rico 00681; Department of Statistics and Data Science, Cornell University, Ithaca, NY 14850, U.S.A.

## Abstract

Single-molecule Förster resonance energy transfer experiments have added a great deal to the understanding of conformational states of biologically important molecules. While great progress has been made in studying structural dynamics of biomolecular systems, much is still unknown for systems with conformational heterogeneity particularly those with high flexibility. For instance, with currently available techniques, it is difficult to work with intrinsically disordered proteins, particularly when freely diffusing smFRET experiments are used. Simulated smFRET data allows for the control of the underlying process that generates the data to examine if a given smFRET data analysis technique can detect these underlying differences. Here, we include a distribution of inter-dye distances generated using Langevin dynamics to simulated freely-diffusing smFRET timestamp data in order to model proteins with conformational flexibility within a given state. We compare standard analysis techniques for smFRET data to validate the new module relative to the base PyBroMo software and observe qualitative agreement in the results of standard analysis for the two timestamp generation methods. The Langevin dynamics module provides a framework for generating timestamp data with a known underlying heterogeneity of inter-dye distances that will be necessary for the development of new analysis techniques that study flexible proteins or other biomolecular systems.

## Introduction

Structure and dynamics of proteins and other biomolecules are fundamental to their function^1^. Static structural data from high-resolution structure determination techniques such as xray crystallography and cryogenic electron microscopy can provide a detailed picture of these systems but they lack information on the transitions between the states^2^. Other experimental techniques may allow for quantitative characterization of the transitions between states^3^, however, but with a reduced spatial resolution. One such technique is single-molecule Förster resonance energy transfer (FRET) spectroscopy or smFRET^3^. FRET is the non-radiative transfer of energy initially absorbed by a "donor" dye to a nearby "acceptor" dye^4,5^. The energy transferred between a donor and acceptor dye is dependent on the distance between the dyes and can be used to provide information on the molecular conformation at the time of photon emission. Therefore, FRET is often considered as a "spectroscopic ruler".^6^ Ensemble FRET experiments, with simultaneous excitation of multiple donors at the same time, contain distance information but they suffer from ensemble averaging that can obscure the protein conformational dynamics underlying the process. Through clever experimental design, valuable conformational information can still be gleaned^7–9^.

The advent of single molecule spectroscopic techniques transformed biophysics into a source of dynamic data on molecular structure as well as function^10^. SmFRET experiments avoid ensemble averaging by taking advantage of exciting the donors and detecting the donor and acceptor signals at a single molecule level^11,12^. These techniques have become a popular source of spatio-temporal information on the conformational landscape of a molecule and have been applied in studies of a variety of systems from DNA^13^, and RNA^14–16^, to protein folding^17,18^.

The two broad varieties of smFRET experiments are distinguished by how the labeled molecule is isolated from other FRET signals when it is excited. First, surface immobilized experiments fix the labeled molecule to a substrate, expose it to laser light to excite the donor dye, and collect the resulting photon data using integrated cameras.^19^ This experimental procedure uses long exposure times to collect data because photon intensities are calculated by integration over multiple pixels where molecules are identified. This setup allows for multiple molecules to be analysed simultaneously but is limited by the frame rate of the camera, which is 1 ms or longer. Despite experimental difficulties arising from surface impacts on dynamics and signal issues from photo-bleaching or other noise sources, surface immobilized experiments have been a fruitful area of study.

Second, freely-diffusing smFRET methods record photon emission timestamps from labeled molecules as they diffuse through a solution with a confocal laser spot focused inside the solution. Periodically, the path of a molecule will cross the focal region of the laser, where the probability of photon absorption and emission are high. The diffusion rates and concentrations of the molecules in solution as well as the size of the focal region are selected so that the observation of simultaneous excitations of more than one molecule is vanishingly rare within a particular observation time window. The molecules emit bursts of photons as they diffuse through focal beam of the confocal laser spot. Optical filters chosen to correspond to the donor or acceptor photon wavelengths isolate the signal into the two detector channels where photon detectors record arrival times with much greater time resolution than the cameras used in the immobilized experimental setups. Simultaneously, background sources of photons are also reported by the photon detectors. Freely diffusing sm-FRET experiments can capture dynamics occurring on faster scales^3^ because the photon detectors record with a finer time resolution compared with the cameras used in surface immobilized experiments. Also, the potential impacts of the surface on conformational dynamics are avoided^20–22^, but the short bursts of data provide a challenge for analysis. Sliding window methods^23^ have been developed to identify the bursts of photons in freely-diffusing timestamp data, which is where the information from the molecule is contained. On the other hand, a Bayesian method for estimating the mean fluorescence from time-correlated, single-photon-counting data has been developed and shown to be useful.^24^ Recent research has effectively identified and quantified the within-burst dynamics in single-labeled single-molecule fluorescence lifetime experiments, shedding light on the complex processes occurring at the level of a single molecule.^25^ Safar et al. introduce a Bayesian non-parametric framework for analyzing single-photon smFRET data under pulsed illumination. The method effectively captures system state transitions and photophysical rates while handling uncertainty from various noise sources.^26^ Many other studies also demonstrated accurate estimations of experimental and simulated data, offering a promising approach for unraveling complex molecular interactions in single-molecule studies.^27–29^ Currently, we also have conventional time-correlated single photon counting, which can detect the time-stamping photon arrival times with ps accuracy and is based on the measurement of the detection timings of individual emitted photons with regard to the time of their excitation.^30^

While sophisticated statistical methodology is essential to the analysis of any smFRET experiment, the literature on this topic has primarily focused on surface-immobilized smFRET^31^. These techniques include histograms^2^, Gaussian mixture models^32^, hidden Markov models (HMM)^13,33–35^, and Bayesian non-parametric approaches^36,37^. The freely-diffusing smFRET technique is gaining in popularity due to its simpler experimental methodology with no need for surface immobilization^38^ and the high time resolution afforded by photon detectors. To further advance the developing fields of smFRET analysis, the ability to realistically simulate the underlying molecular processes in a systematic, controlled, and reproducible manner is a necessity.

Several existing smFRET software tools have been developed to generate realistic simulated data^39^. Some of these tools generate binned photon data with specific distribution characteristics^40^. Other software, like Fretica^41^ or simFCS 4,^42^ simulates the diffusion of molecules through solution as well as the emission of photons from fluorescent dyes.

In reality, the distances between the dyes on a labeled molecule (dye-dye distance) are constantly changing due to the thermal fluctuations of the molecule. A fixed efficiency assumes that the heterogeneity of dye-dye distances from molecular motion in a freely-diffusing molecule is negligible compared to the other parts of the simulation. This simplification may be justifiable for highly structured molecules or at low temperatures. Reductions in molecular structures and greater fluctuations, like those observed in disordered proteins^43,44^, will invalidate the assumption. This is especially true for disordered proteins with reduced secondary and tertiary structure to stabilize the conformations. The flexibility of the molecule leads to a heterogeneous conformational ensemble that poses further challenges to the analysis of experimental data. Ideas from polymer dynamics have been applied to the describe the smFRET data from disordered proteins^45,46^. Biologically important systems often contain large heterogeneity of conformational states^47^, so analysis of disordered protein systems has great applicability for future work.

To more accurately model the conformational heterogeneity of dye-dye distances of a flexible molecule during an smFRET simulation, an overdamped Langevin dynamics simulation was integrated into freely-diffusing smFRET timestamp simulation data to model the internal conformational dynamics of the molecule. The Langevin dynamics will produce a trajectory of dye-dye distances for each molecule that conform to an underlying ground truth related to the free energy used in the Langevin dynamics. This addition provides a more realistic smFRET simulation, particularly important for unstructured proteins or those associated with intrinsic disorder. This added realism will be necessary in the development of new analysis techniques that account for conformational heterogeneity.

The remaining sections of this paper will describe the simulation methods for generating simulated freely-diffusing smFRET timestamps, including a description of overdamped Langevin dynamics used to generate the distribution of dye-dye distances. Then, two example simulations are described to generate simulated data; one for molecules in a single state and for molecules that inter-convert between two states. The results of a typical analysis methods applied to the example simulations are then presented. A standard analysis for smFRET data is presented in Results using thresholds and Gaussian mixture models was applied to the timestamp data for the single state data using the base PyBroMo software (non-Langevin) and the extended PyBroMo using the added Langevin dye-dye distances (Langevin). Then, an analysis of the two-state inter-conversion methodology using the non-Langevin and Langevin timestamp data uses a skewed Gaussian mixture model, as well as changepoint analysis and hidden Markov models (HMMs) to assess the dynamics states. Finally, we discuss the results presented and presents conclusions based on the comparison of the analysis for the two simulated data sets.

## Methods

Timestamps are generated using the freely-diffusing smFRET simulation package PyBroMo^48^. Details on how PyBroMo simulates diffusion of molecules and emission of photons are contained in the Supplemental Material. To illustrate the impact of conformational dynamics on analysis results, we simulate timestamps with fixed efficiency states and with conformational heterogeneity using Langevin dynamics.

### Overdamped Langevin Dynamics

The use of a static efficiency in the freely-diffusing smFRET simulation implies either an underlying static relationship between the two fluorescent dyes labeling the molecule or dynamics between the dyes that are undetectable within a photon burst. Fluctuations in molecular structure, particularly in unstructured proteins, could impact how smFRET data is interpreted. To include conformational heterogeneity beyond the static efficiency assumptions, an overdamped Langevin dynamics simulation is added to generate realistic dye-dye distance fluctuations over the simulation time as a one dimensional diffusion process within a free energy field.

The Langevin trajectories are calculated according to the Euler-Maruyama method^49^, where at each time step (δ t), the dye-dye distance is updated by calculating the contributions from the distance derivative of the free energy function, *V* (*r*) and a stochastic random contribution. This step update is defined as

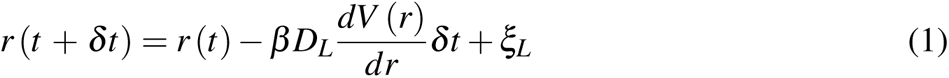

where *D_L_* is the diffusion coefficient, 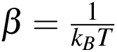 with *k_B_* being the Boltzmann constant, *T* is the system temperature, and ξ*_L_* is drawn from a Gaussian distribution of mean 0 and variance 2*D_L_*δ *t* (ξ*_L_ ∼ N*(0, 2*D_L_*δ *t*)). The diffusion coefficient for the dye-dye distance, *D_L_*, is distinct from the Brownian motion diffusion coefficient. The *D_L_* here is the diffusion coefficient associated with the conformational dynamics of protein, which not only depends on the specific protein but also, for the same protein, depends on the location of the attachment of the dyes. In some cases, the inter-dye distance may fluctuate very fast not visible to the smFRET technique (i.e., fluctuations are much faster than 1 ms). In other cases, the inter-dye distance may change on a second or minute timescale. The molecules are perturbed by thermal white noise while inside a user defined free energy field.

A FRET efficiency model converts the dye-dye distance trajectories to efficiency trajectories. Two different efficiency models are used for the two example scenarios described in greater detail in the Methods section. However, a constant that is common in efficiency models is the Förster radius, *R*_0_, defined as the distance from the donor dye at which FRET efficiency is 0.5. This *R*_0_ value is specific to the fluorescent dyes used in a smFRET experiment and based on the quantum yield of the donor dye and the spectral overlap of the two dyes. The photon emission simulation takes the efficiency trajectories and uses the efficiency value at the time of photon emission and uses it to determine the ratio of donor and accetor photons. The background timestamp generation is independent from the Langevin dynamics and contributes to the Poisson distributed background timesteps as before.

Next, we describe the two example simulations to compare simulated data with and without conformational heterogeneity from Langevin dynamics.

### Molecules in a Single State

To demonstrate the generation of timestamps using the Langevin dynamics conformational trajectories, a simple example system of molecules in a harmonic free energy field is simulated for three independent simulations with all parameters held constant. A Gaussian distribution of distance is a common assumption in polymer physics, though other distributions have been explored to account for more complex system interactions^50–52^. The harmonic function, *V_H_* is defined as

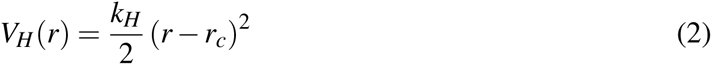

where *k_H_* is the harmonic force constant, and *r_c_*is the center of the harmonic well. 100 molecules are contained in a simulation box with lengths *L_x_*= *L_y_* = 8 µm, *L_z_* = 12 µm. The Brownian diffusion coefficient, *D_B_*, is set to 30 µm^2^/s for all molecules. Diffusion coefficients for proteins in water are on the order of 20 *−* 200 µm*/*s^53,54^ making this value on the slower side of the spectrum. A faster diffusion rate would decrease the average duration of a burst of photons as the molecule traversed the focal spot in a shorter period of time. The Gaussian point spread function (PSF) is centered in the simulation box with a σ*_x_* = σ*_y_*= 0.3 µm, and σ*_z_* = 0.5 µm. Three independent simulations are run for 10s each with a time step, δ *t*, of 50 ns. For timestamp generation, a maximum emission rate of 200,000 counts per second (cps) is used in all the simulations, as well as a background rate of 1,200 cps for the acceptor channel and 1,800 cps for the donor channel. The cps values will be kept consistent for all simulations used in this work.

For the Langevin dynamics parameters, the thermodynamic coefficient β is 1.339 (kcal/mol)*^−^*^1^ which corresponds to a relatively high temperature of 378 K for large thermal fluctuations. The Langevin diffusion coefficient, *D_L_*, is 1300 Å^2^/ns. This is a rapid diffusion to explore the free energy well in a short trajectory. The harmonic function is defined by Eq. (2) with the coefficient *k_H_* set at 0.025 (kcal/(mol Å^2^)) with the center of the harmonic well at 40 Å for 50 of the molecules, and at 65 Å for the 50 molecules. Eq. (4) is used to convert the distances to efficiencies. In efficiency conversions, an *R*_0_ of 56 Å is used.

A short trajectory of Langevin dye-dye distances is shown in Figure 1. The molecules in each population oscillate in the harmonic free energy field over time, with a probability of some dye-dye distance, *P*(*r*), following the relation

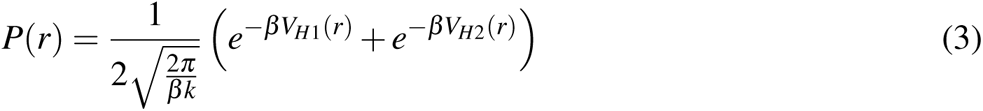

**Figure 1:**
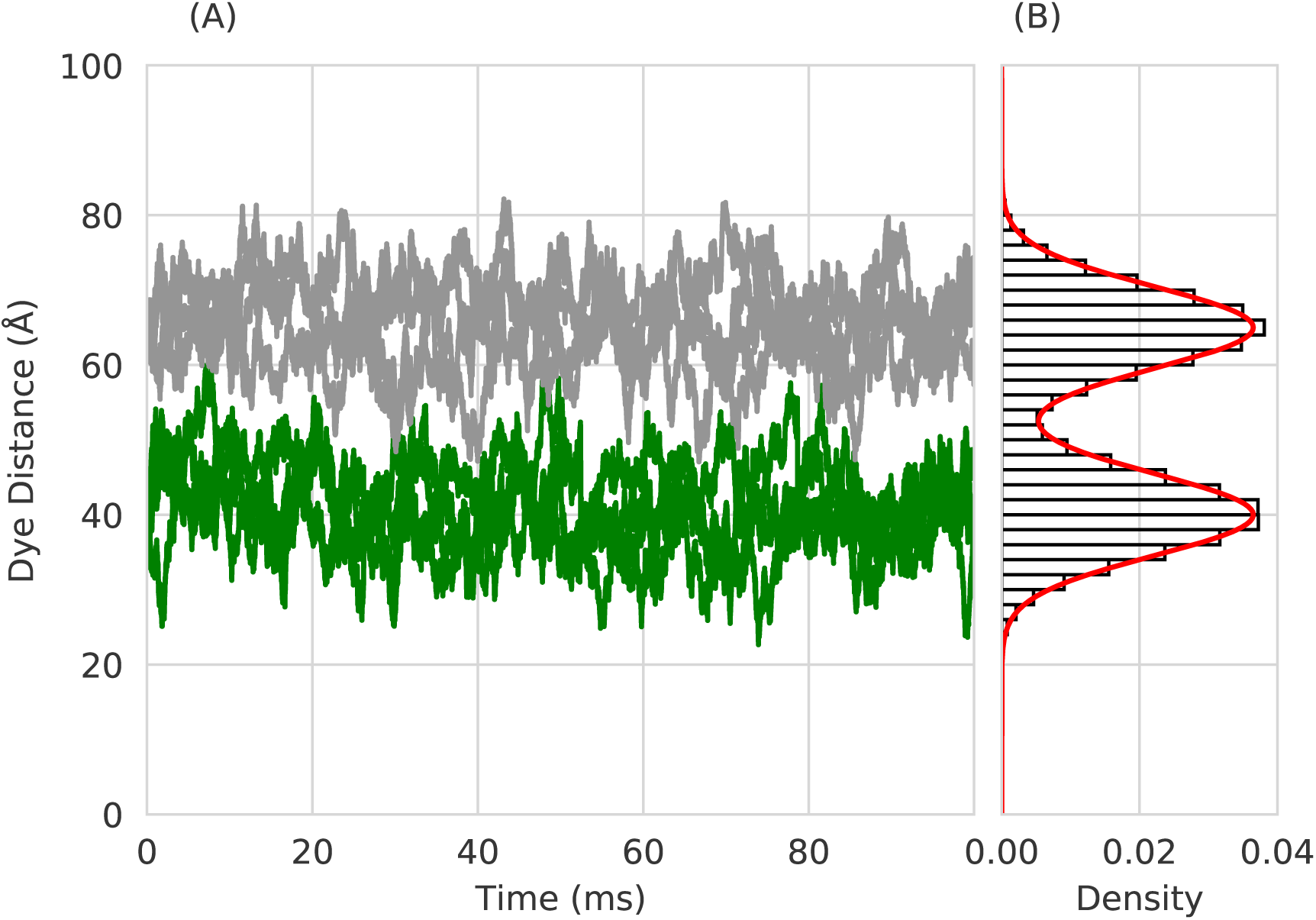
(A) A portion of a trajectory of Langevin dynamics for the dye-dye distance of 4 molecules in a harmonic free energy centered at 40 Å (colored green) and 4 molecules in a harmonic free energy centered at 65 Å (colored gray). (B) A histogram of the dye-dye distances for the combined populations along with the analytic solution for the distribution in red.

where *V_H_*_1_ and *V_H_*_2_ are the harmonic free energy functions applied to the two molecule populations in the Langevin dynamics simulation.

For the harmonic free energy simulations, an efficiency model developed for conformationally heterogeneous proteins^44^ relating the dye-dye distances to FRET efficiency is,

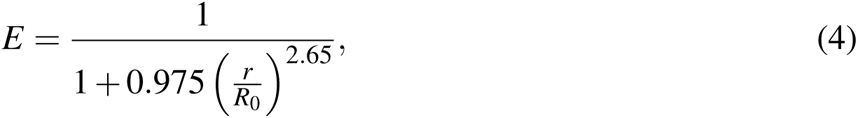

where *r* is the dye-dye distance and *R*_0_ was 56 Å. This efficiency model is used in part because this work focuses on the heterogeneous conformations of proteins, but more to demonstrate general approach to efficiency that this simulation uses. In the next example, the more standard inverse 6*^th^* power efficiency model is used. The FRET efficiencies used for photon generation are 0.41 and 0.71, which corresponded to Eq. (4) applied to distances of 40 Å and 65 Å, respectively. The distances matched the centers of the harmonic functions used in the Langevin dynamics simulations. The other photon generation parameters for maximum emission rate and background noise were held constant.

To compare the results of the simulations with the new Langevin dye-dye distance trajectories with the fixed efficiency assumption, three sets of simulated timestamps were generated with the base (non-Langevin) PyBroMo. These simulations used the same number of molecules, and other Brownian motion parameters for diffusion coefficient, simulation box, PSF, and background photons as described above. 50 molecules had an efficiency of *E* = 0.71 while the other 50 had an efficiency of *E* = 0.41. These efficiency values correspond to Eq. (4) applied to the harmonic centers from the Langevin dynamics, 40 Å and 65 Å respectively.

### Molecules with Inter-conversion Between Two States

The harmonic free energy Langevin simulations described above approximate a system where the dye-dye distance fluctuates around a single state for the duration of the simulation. However, biophysical intuition as well as experimental smFRET data suggest that many biomolecular systems correspond to two or more interconverting conformational states at equilibrium^55^.

To simulate a system that dynamically moves between different states, a bistable free energy with two symmetric wells are applied to a system of molecules in the Langevin dynamics simulation. 90 molecules were simulated using the Langevin dynamics simulation with the same *D_B_*, time step length, simulation box, and PSF as previously defined in single state molecule simulations. This bistable function, *V_B_*(*r*), is defined as

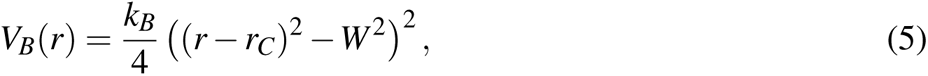

where *k_B_* is the bistable force constant set at 10*^−^*^4^ (kcal/(mol Å^2^)). The location of the center of the bistable function, *r_C_*, is set in this example at 50 Å, and *W* is the the offset from the center where the free energy wells were located, set at 15 Å. The locations of the bistable minima is at *r_c_ ±W*, or 35 and 65 Å with a barrier height of 1.265625 kcal/mol. Using the bistable free energy function, a Langevin molecule will explore a local free energy energy well until a large enough energetic contribution from the white noise in the Langevin dynamics gives the molecule the energy to overcome the energy barrier and explore the other well. A Langevin diffusion coefficient of *D_L_* = 0.002 Å^2^/ns is used. This value is selected for slower dynamics.

FRET efficiency is modeled using the commonly used relation

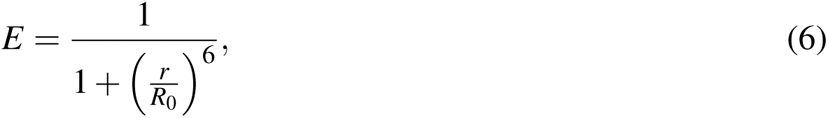

where *r* is the dye-dye distance and *R*_0_ = 56 Å, as before. The efficiency model in Eq. (6) is based on a point dipole-dipole approximation and is widely used in smFRET data analysis.^19^ The dye rotational dynamics is assumed to be considerably faster than protein conformational dynamics. In order to gather a sufficient amount of data for analysis, a total of approximately 20 minutes of simulated smFRET data is generated.

A short dye-dye distance trajectory using the bistable function is shown in Figure 2. We see the dye-dye distances oscillate inside one of the free energy wells for some period of time before eventually overcoming the energy barrier between the two wells and switching states. The distribution of dye-dye distances for the bistable Langevin simulation follows the relation

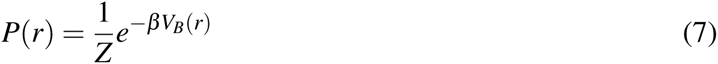

**Figure 2:**
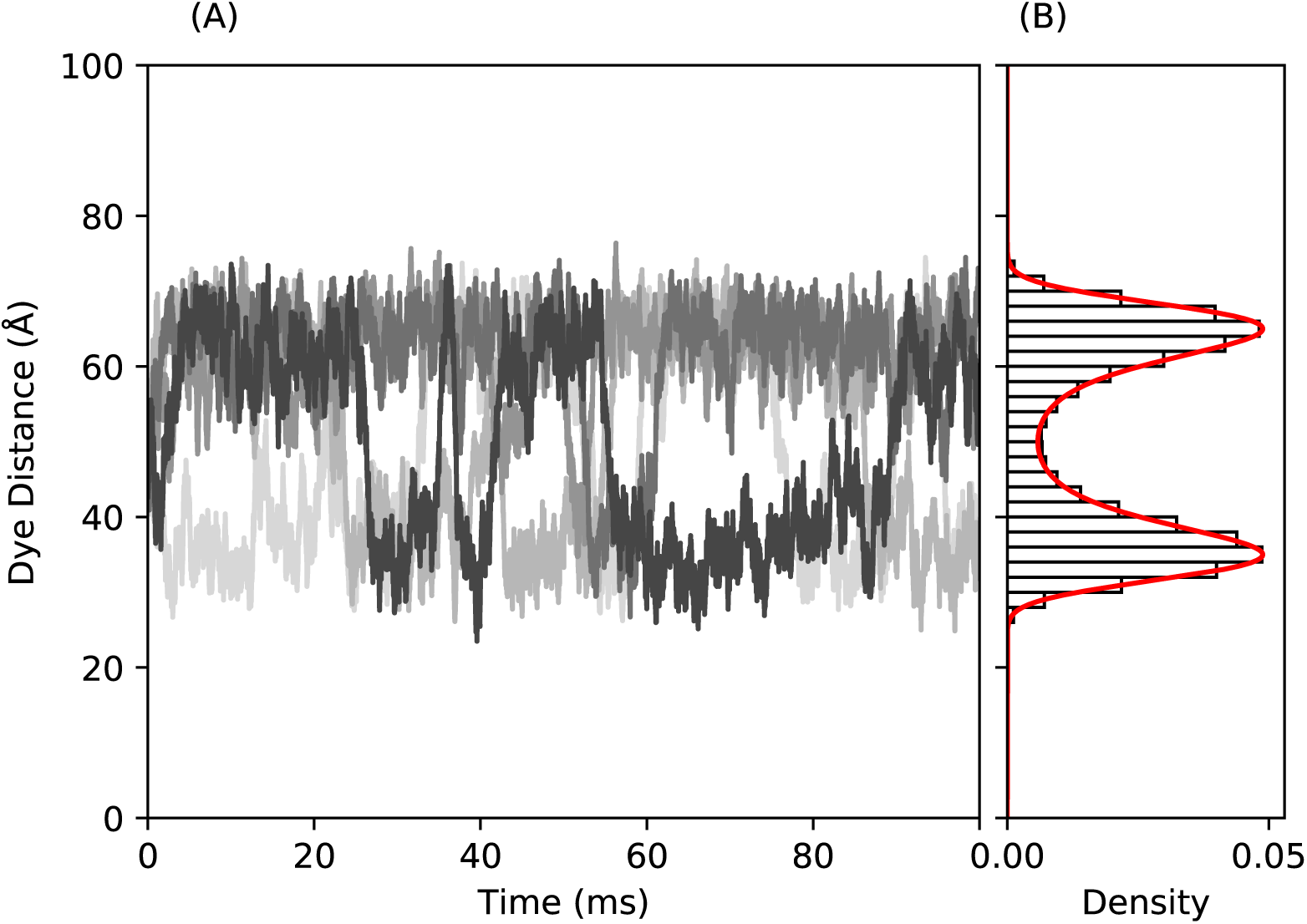
(A) A portion of Langevin dynamics trajectories for the dye-dye distance of 5 molecules moving in a bistable free energy field centered at 50 Å with minima at 65 Å and 35 Å. (B) A histogram of the the dye-dye distances is shown with the theoretical probability shown as a red line.

where the partition function for the bistable free energy, 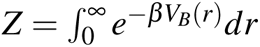, normalizes the probability density to 1. A lower temperature, *T* = 300K, is used as compared to Example 1, with β = 1.679 (kcal/mol)*^−^*^1^. The lower temperatures decrease the magnitude of thermal fluctuations for each time step so the molecule will explore the local well long enough to emit sufficient photons for the state to be identifiable.

The analytical transition matrix of the bistable Langevin simulation, *T* (0), between different states is related to the transition rate matrix, *Q*, by

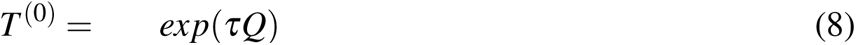

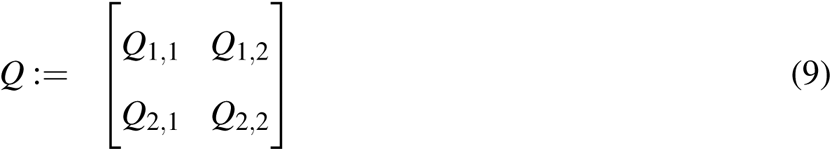

where τ is the lag time between state determination measurements. The entry *Q_i, j_*represents the transition rate from state *i* to state *j*. The transition rate between two non-identical states (here reactant, *R* and product, *P*) is calculated using relations from Berezhkovskii and Szabo,^56^

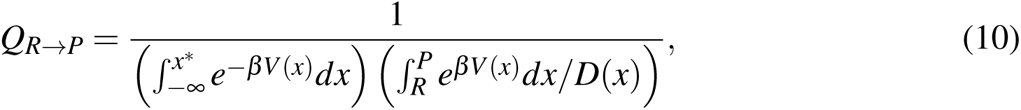

where the integration limit *x^∗^* is the peak of the barrier at 50 Å, *V* (*x*) is the free energy, *D*(*x*) is the position dependent diffusion coefficient, and 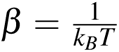. Substituting in the bistable function, *V* (*x*) = *V_B_*(*x*) and the constant Langevin diffusion coefficient, *D*(*x*) = *D_L_*, the transition matrix can be computed theoretically as,

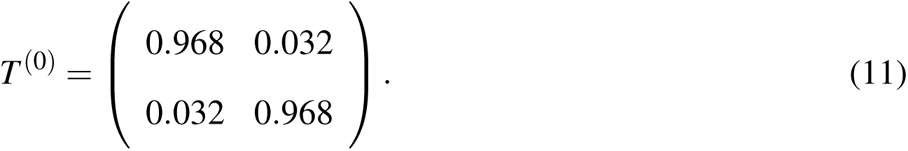

In addition to the 90 molecules in the bistable Langevin simulation, 10 molecules were kept in a constant "donor only" state of *E* = 0. Donor only states are present in experimental data and represent molecules where only the donor dye is attached, with no FRET possible. The donor only population represents a phenomenon of imperfect labeling where the acceptor dye is not present as this is a source of error encountered by experimenters. This is a static state and does not adequately describe other similar error sources like photoblinking, causing a temporary donor only signal.

To provide a comparison with the bistable Langevin timestamps, non-Langevin timestamps were generated that simulated dynamic state switching. This is done by generating two timestamp traces of approximately 20 minutes in length, using the same parameters for Brownian motion as the bistable Langevin data. One set of timestamps used a fixed high efficiency state of *E* = 0.944, while the other used a fixed low efficiency state of *E* = 0.290. The efficiencies correspond to Eq. (6) using the locations of the well minima, *r_C_* = 35 Å for the high efficiency state and *r_C_* = 65 Å for the low efficiency state Also, a Förster radius of *R*_0_ = 56 Å was applied in all the efficiency calculations. Again, the Brownian motion simulations parameters of Brownian diffusion constant, simulation box size, PSF, and background photons were the same for the non-Langevin simulation as with the Langevin dynamics simulations above.

Transitions between states were simulated by drawing residence times from an exponential distribution with an average residence time of 31.126 ms. The trajectory of an efficiency state evolves like a step function alternating between the two states. This residence time leads to a transition matrix for the non-Langevin data that closely matches the transition rate matrix generated from the bistable free energy. Using these residence times, a set of timestamps is created that switched between the two efficiency states, also 20 minutes in overall length.

A comparison of the results from three analysis methods performed on the dynamic state model simulation using non-Langevin and Langevin timestamps are contained below.

## Results and discussion

Techniques for simulating freely diffusing smFRET experiments are valuable, in large part, because they allow researchers to evaluate statistical methods using realistic data with a known ground truth. With this in mind, we present a standard analysis of the timestamp data produced from the parameters described in the Methods section.

### Comparison of Single State Simulations

The timestamp data generated by both non-Langevin and Langevin simulations was in the form of a column of ordered timestamps when a photon was detected. Additional columns label the channel that detected the photon (donor or acceptor), and a label to identify the molecule that emitted the photon. This molecule identifier would not be available in experimental data, but is information that is available in the simulation.

Data analyses of freely diffusing smFRET experiments typically begin by binning and thresholding the raw photon time stamp data^31,57^. The time bin size needs to be long enough to collect sufficient data such that the signal from the fluorescent dyes can be distinguished from the noise contributions. Conversely, the bin size needs to be small enough so that the FRET signal is only from one molecule. The specific choice of time bin length will be dependent on background noise rates, molecule diffusion rates, and confocal beam size, on the order of 1 ms.^23^ In our analyses, we use a typical experimental bin width of one millisecond. For a given experiment, let 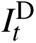 and 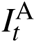 denote the photon counts in the donor and acceptor channels during time bin *t* and define the combined count 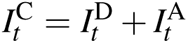. We restrict our analyses to those time bins with combined count exceeding 40 photons. Based on the simulation parameters that are used, a combined photon count at or above this magnitude indicates that the signal is very likely from a molecule diffusing across the focal beam and thus the proportion of photons in the acceptor channel reflects the molecule’s conformational state. Threshold also ensures that our estimates of the efficiencies within each time bin are not excessively variable due to low counts. No single method to determine photon thresholds has been universally accepted^58^. In the literature, there are a number of heuristics for choosing the threshold and many alternative approaches to identifying the diffusion of a molecule across the focal beam^59–61^.

Central to our analysis are the estimates of efficiencies within each bin, which we refer to as *apparent efficiencies*.^19^ The apparent efficiency within bin *t* is defined as the proportion of the total photon count from that bin which was detected in the acceptor channel:

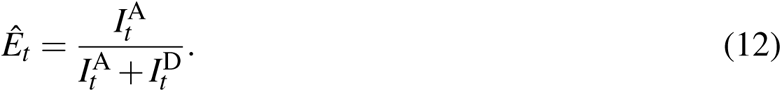

When analyzing real smFRET experiments, estimation of efficiencies should also take into account the so-called γ factor, which accounts for the difference in quantum yields of the donor and acceptor dyes as well as the difference in photon detection efficiencies of the donor and acceptor channels.^62,63^ Other experimental error sources, include adjustments for direct excitation of the acceptor dye from laser and leakage of acceptor photons into the donor channel^64^. These adjustment is not necessary for our analysis because the smFRET simulations in this article were run with equivalent quantum yields and equivalent detection efficiencies.

We analyze the simulated smFRET experiments using a simple histogram of the apparent efficiencies as well as a Gaussian mixture model fit to the apparent efficiencies. The histogram approximates the marginal distribution of efficiencies. It provides an idea of the relative amount of time a molecule spends at each efficiency and whether there exist easily-distinguished conformational states. In comparison to a histogram-based analysis, the analysis based on a Gaussian mixture model provides more quantitative information related to hypothesized latent conformational states. We suppose that there is a latent conformational state *s_t_ ∈ {*1*,…, K}* associated with each time bin *t* and that these latent conformational states are independent and identically distributed with probabilities π_1_*, …,* π*_K_.* Given that *s_t_* = *k,* we suppose that the apparent efficiency 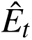 follows a Gaussian distribution with mean µ*_k_* and variance 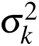. The smFRET simulations were run with *K* = 2 conformational states, and we take this as given. We compute the maximum likelihood estimates of the unknown parameters via an expectation-maximization algorithm^65^ as implemented in the mixtools package^66^ in R^67^.

$$Figure 3 compares the non-Langevin and Langevin simulations in terms of apparent efficiencies and the corresponding dye-dye distances. Figure 3 (A), based on the non-Langevin simulation, shows the estimated two-component Gaussian mixture density (in solid black) on top of a histogram of the apparent efficiencies. The dashed lines represent the (weighted) densities of the estimated component distributions. The low efficiency component has a mean of 0.42, a standard deviation of 0.07, and a mixture weight of 0.62. The high efficiency component has a mean of 0.70, a standard deviation of 0.05, and a mixture weight of 0.38. The vertical red arrows are placed at the true efficiency values used in the simulation. Figure 3 (B) shows the corresponding histogram, densities, and arrows after a transformation to the distance space. The probability distribution of distances is converted to a probability distribution of efficiencies through a change of variable based on the efficiency model in Eq. (4) The conversion is over the apparent efficiencies which represent an average over 1 ms of data. This averaging has the potential to obscure faster internal dynamics that occur within 1 ms time bin.

**Figure 3:**
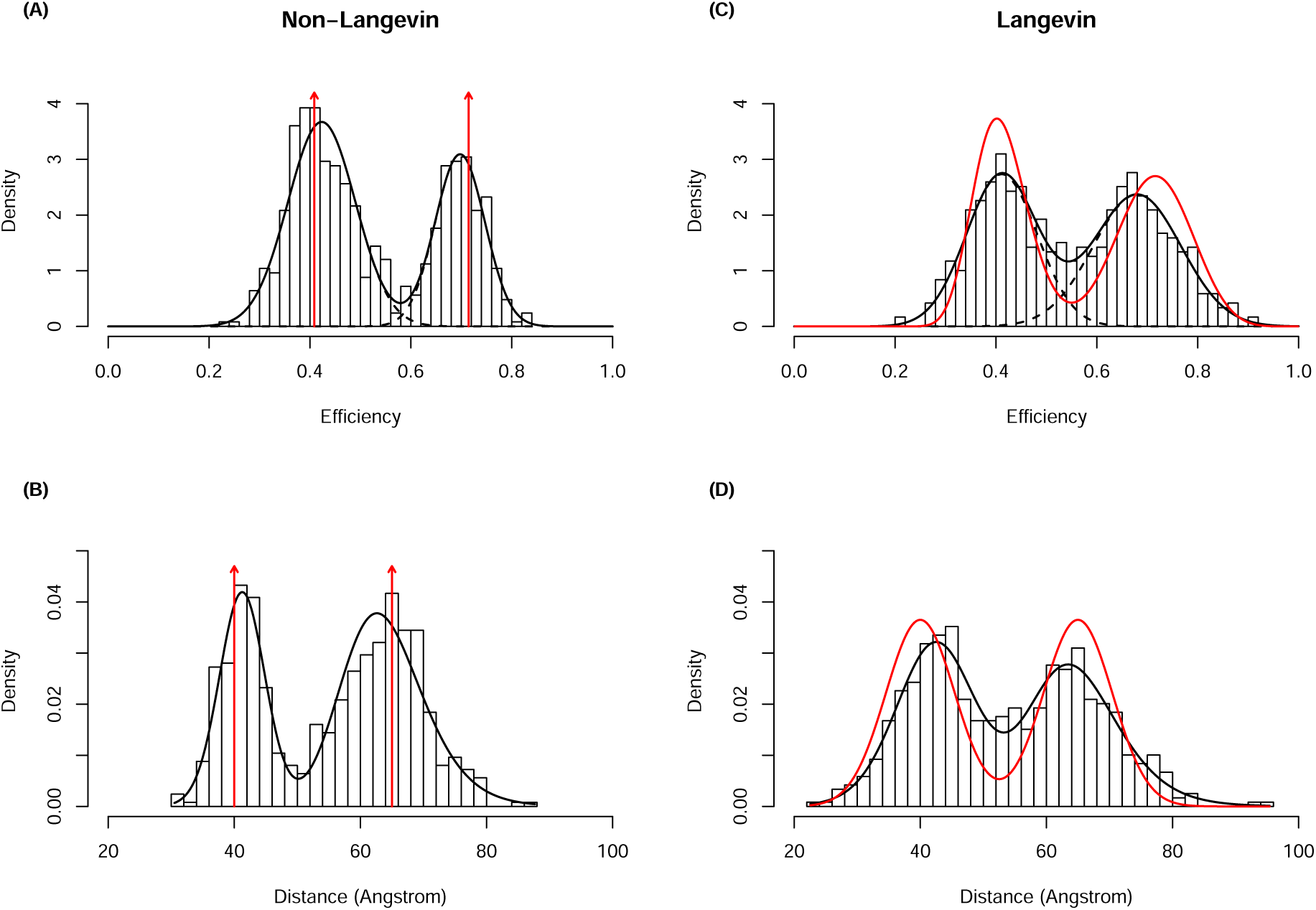
(A) The estimated Gaussian mixture density (solid black line) from the non-Langevin simulation on top of a histogram of the apparent efficiencies along with the two true efficiencies (red vertical arrows). (B) The corresponding plot in the distance space. (C) The analogous efficiency plot for the Langevin simulation. (D) The analogous distance plot for the Langevin simulation. The density curves in red shown in C & D are the "ground-truth" analytical distributions of efficiency and distance, respectively, associated with the Langevin model. The vertical arrows in A & B are the corresponding distributions in the non-Langevin model that represent Dirac delta functions.

In contrast to Figures 3 (A) and 3 (B), which are generated from the Non-Langevin simulation, Figures 3 (C) and 3 (D) are based on the Langevin simulation. For a better comprehension of the graphs, Langevin simulation results 3 (C-D) contain density curves showing the true, non-degenerate theoretical distribution of efficiencies (or distances) in place of vertical red arrows at two actual efficiencies (or distances) in 3 (A-B). The two-component Gaussian mixture as specified by Eq. (3) is the theoretical distribution in the distance space. Again, using the same equation, Eq. (3), and changing the variables from efficiency to distance, we calculated the theoretical distribution in the space of efficiency. The low-efficiency component in Figure 3 (C) has a mean of 0.41, a standard deviation of 0.07, and a mixture weight of 0.48, whereas the high-efficiency component has a mean of 0.68, a standard deviation of 0.09, and a mixture weight of 0.52. Figure 3 displays the underlying Langevin dye-dye distance distribution as a red line Figure 3 (C-D). Compared to the Langevin simulation timestamp analysis, which exhibited a larger distribution with more overlap, the separate peaks shown in the non-Langevin timestamp analysis revealed less overlap in the distribution of the two populations. Due to the overlap between the underlying distance distributions of the Langevin dynamics for the two populations, this little discrepancy is understandable. Overall, the research demonstrates that the addition of over-damped Langevin dynamics to a straightforward situation results in timestamps that include important data from the underlying distance distribution, such as the locations of efficiency peaks. We note that we do not expect the analytical distributions and the simulation data to be identical since what we describe as the analytical distribution here is simply based on the conformational dynamics of a fixed protein in a theoretically idealized case, where both the Brownian dynamics and the shot noise process are both ignored. As a result, even in the case of the non-Langevin model, we expect the broadening of the observed distributions although the analytical distributions are Delta functions.

### Comparison of State Inter-conversion Simulations

Next, we describe two more sophisticated analyses that account for additional realistic features included in the simulation, like donor-only particles and dynamic state changes. A histogram based analysis as well as analyses to infer state dynamics were performed. Again, the non-Langevin and Langevin timestamps generated used the simulation parameters that contained information consistent with the simulation parameters that was detectable by the analyses.

#### Skewed-Gaussian Mixture Model

We again analyze the non-Langevin and Langevin timestamps through mixture models. This time, we fit three component skewed-Gaussian mixture models to the timestamps generated from Example 2. Adding a third component is necessary because these simulations include donor-only molecules, leading to a low FRET peak. The skewed-Gaussian distribution has density

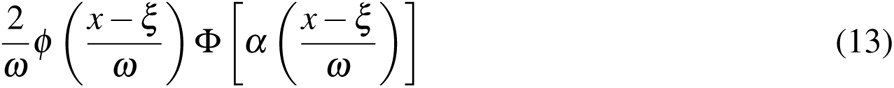

where ϕ and Φ are the density and distribution functions of a standard Gaussian random variable, ξ is a location parameter, ω is a scale parameter, and α is a shape parameter^68,69^. This more flexible parametric family allows us to adequately model skewed distributions. Apparent efficiency distributions which lie near the boundary of the unit interval, including the low FRET peak, typically exhibit strong skewness. We compute the maximum likelihood estimates of the unknown parameters via an expectation-maximization algorithm as implemented in the mixsmsn package^70^.

The results appear in Figure 4, which compares the non-Langevin and Langevin simulations in terms of apparent efficiencies and the corresponding dye-dye distances. Figure 4 is analogous to

**Figure 4:**
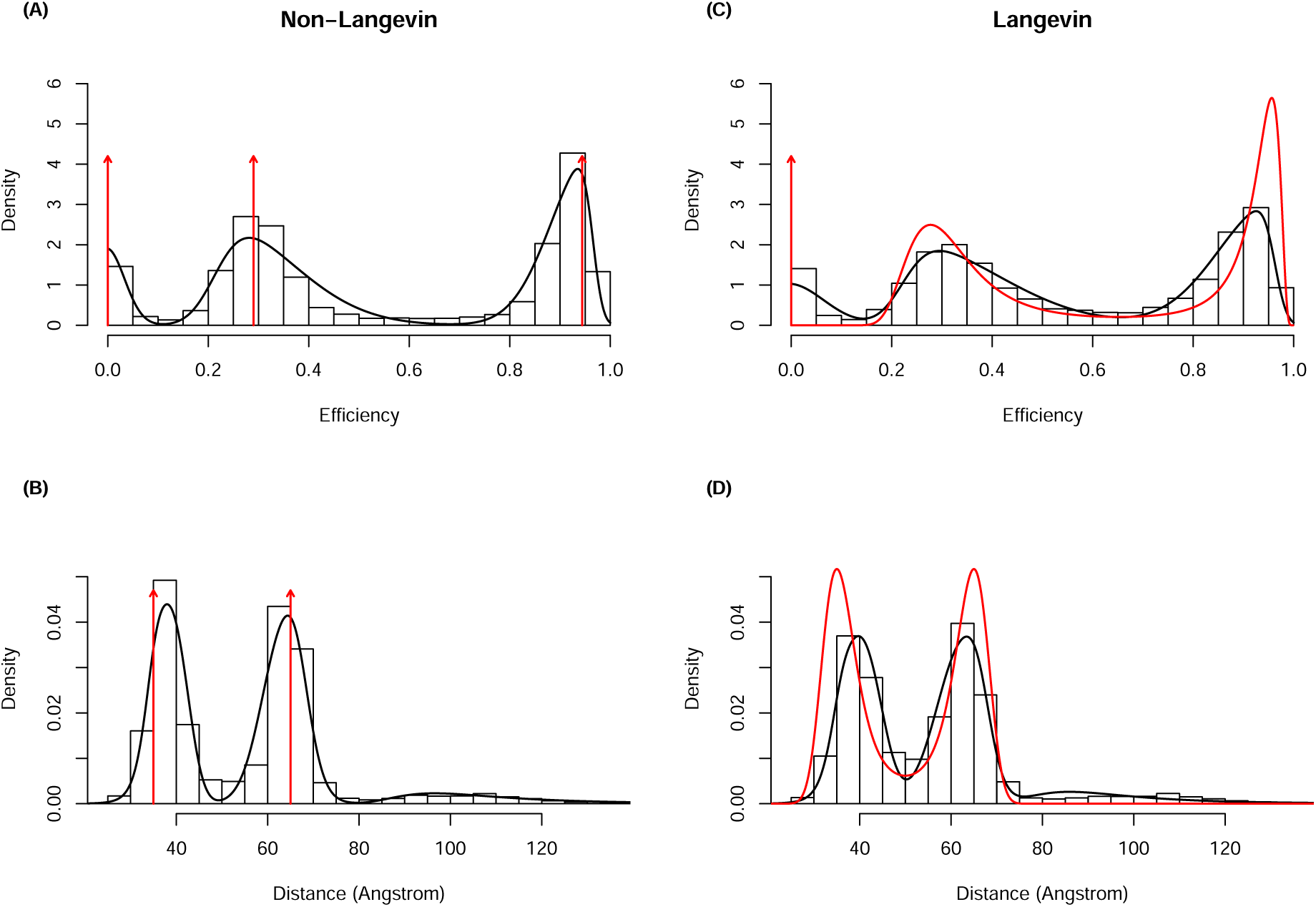
(A) The estimated skewed-Gaussian mixture density (solid black line) from the non-Langevin simulation on top of a histogram of the apparent efficiencies along with the true efficiencies (red vertical arrows). (B) The corresponding plot in the distance space. (C) The analogous efficiency plot for the Langevin simulation. (D) The analogous distance plot for the Langevin simulation. The density curves shown in red in C & D are the true, non-degenerate theoretical distribution of efficiencies (or distances).

Figure 3, except here they depict the results of the skewed-Gaussian mixture model. The skewed Guassian mixture analysis was able to recover the location of efficiency peaks from the timestamp data reasonably well for both the non-Langevin and Langevin data, as well as the donor-only peak. Again, the efficiency states for the non-Langevin simulation timestamps showed higher, more well defined peaks with less overlap than the Langevin simulation timestamps, consistent with the point mass distribution in the distance. This method aggregates all the timestamp information over time into a histogram, losing temporal information about switches between states. The next two analysis methods will explore the state switching in the timestamp data with more depth.

#### Burst Analysis

In the previous histogram analyses, a threshold placed on the binned timestamp data was used to identify the bins with FRET signal "bursts" that occur when a molecule diffuses through the focal spot and background noise. More sophisticated methods have also been developed to identify bursts of FRET signal in timestamp data.

FRETBursts^58^ is used to identify bursts with the simulated data from example 2. This software applies the "sliding window" search algorithm^60^. For our initial search, we applied a window of 200 consecutive timestamps with a threshold of 40,000 cps to that window. After the initial search, only bursts with at least 100 photons are selected. The selection of parameters for a burst search can be somewhat arbitrary, though the general approach was to pick a threshold higher than background. Some paramaterizations will estimate the background from the data and then apply a threshold based on the estimated background.

A burst variance analysis^71^ (BVA) using FRETBursts is done to compare the non-Langevin and Langevin timestamp data. Each burst was split into sub-bursts of 10 photons in order to calculate the standard deviations, σ*_i_*efficiencies in Figure 5. A 2D histogram with a kernel density estimate (KDE) smoothing presents the smooth contour plots of the distribution of efficiencies and σ*_i_*. A dashed line represents the σ of a binomial distribution for the size of the sub-burst. State peaks that occur on the dashed line are static, while state peaks above the dashed line contain inter-conversion of states within a burst. In both data sets, the two FRET states at 0.29 and 0.94 are visible with the small peak from the donor only contribution visible in the lower left of each plot. Both state peaks in the non-Langevin analysis are centered on the dashed line, indicating negligible dynamic heterogeneity, as expected. Conversely, both state peaks of the Langevin data are near the dashed line, but slightly above, indicative of a small dynamic heterogeneity. More visible differences arise in the areas representing inter-converting states that arc between the main peaks and have higher variances. In the non-Langevin data, the inter-converting arc is less populated, while the Langevin data shows a higher population for this high-variance region; although it still represents a small population compared to the two main peaks. These observations are consistent with the differences in the underlying process of state inter-conversion for the Langevin and non-Langevin data. In the non-Langevin model, the states inter-convert in discrete states and fewer bursts would capture this interconversion as compared to the Langevin model, where the states inter-convert more gradually. The slight shift in the two main peaks of the Langevin model is also explained with the fact that these states are not associated with fixed distance/efficiencies. In other words, one expects dynamic fluctuations within each of the two major states. A different metric called FRET-two-channel kernel-based density distribution estimator (2CDE)^72^ estimates the density of photons in each channel for each photon in a given burst. Analysis of FRET-2CDE in the Supplemental Materials shows a similar difference between the non-Langevin and Langevin data from Example 2 with the arc of inter-converting states between the main peaks being thicker with higher density of photons in the Langevin data.

**Figure 5:**
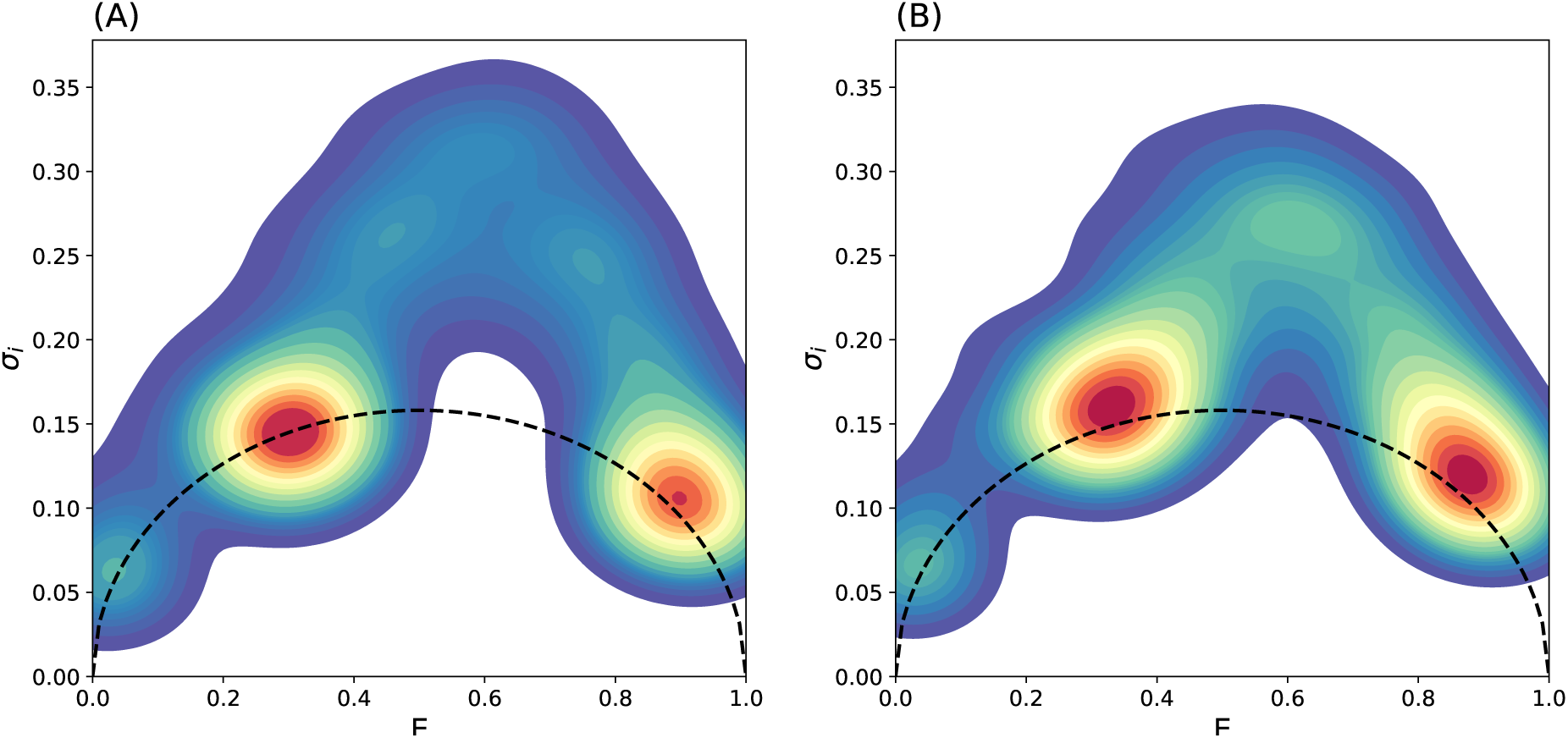
BVA to compare the (A) non-Langevin and (B) Langevin data. Each burst was split into sub-bursts of 10 photons to calculate the standard deviation. A KDE is used to present smooth contours the scatter plot of efficiency and variance of efficiency for each burst. The dotted line shows the standard deviation of a binomial distribution for 10 samples.

#### HMM Analysis

We analyze the Example 2 timestamp data using a hidden Markov model (HMM)^73^. Specifically, we consider only the time-bins which are above a threshold (where the total photon count is above 40). In contrast to surface immobilized smFRET, in freely diffusing smFRET experiments the molecule is only sometimes in front of the focal spot^35,58^. Other researchers have used a burst search algorithm^58^ to identify the parts of the trajectory when the labeled molecule is in the focal spot. A "window search" algorithm^60^ is employed in a burst search analysis. For this analysis, we define a burst region as a set of consecutive time bins such that for each of them, the total photon count is above the threshold. We then evaluate the sequence of apparent efficiencies for each burst region. To perform dynamical analysis and detect transitions between the different FRET states, we treat the sequence of apparent efficiencies from each burst region as an independent time-series to be modeled with the HMM^73,74^, where the HMM parameters are constant for all the independent time-series. We fit the apparent efficiencies using two hidden states, and assume they are normally distributed conditionally on each state. We have restricted the HMM model to two states with distribution analysis above a threshold of 40 photon count from Example 2. Python’s hmmlearn^75^ package was deployed to fit the HMM.

For the data generated using Langevin dynamics, the average photon burst region duration is 2.18 bins of 1*ms*. We fit the HMM using a total of 30053 such burst regions and obtain a transition matrix estimate

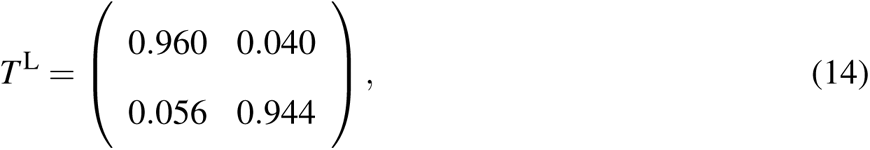

corresponding to two Gaussian states, for which we estimate means, µ_1_ = 0.321, µ_2_= 0.883, and variances 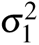= 0.029, 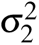= 0.004, respectively.

For comparison, we analyze the data generated using non-Langevin dynamics, where the average photon burst duration is 2.20 bins of 1*ms*. We fit the HMM using a total of 31354 such burst regions. Fitting the data results in a transition matrix:

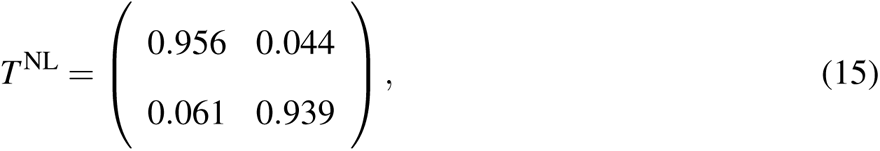

corresponding to two Gaussian states, with means, µ_1_ = 0.291, µ_2_ = 0.910, and variances 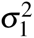= 0.025, 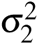= 0.002, respectively.

Qualitatively, the measured transition matrices for both the Langevin and non-Langevin models look reasonably similar to the analytical transition matrix in Eq. (11). The Langevin model, however, results in a transition matrix that is slightly closer to the ground truth. This is verified using a careful quantitative analysis of the difference between the known and measured transition matrices along with error analysis, presented in the Supporting Information.

A visualization of the transitions using changepoint analysis is presented in the Supporting Material, Figure S2, and shows reasonable qualitative agreement between Langevin and non-Langevin simulations. From these results we can infer that the Langevin dynamics module produces timestamps that include dynamic state changes in a controlled and realistic manner.

## Discussion

The new Langevin dynamics allows for generating more realistic smFRET data consistent with what one expects to observe from freely-diffusing smFRET experiments of molecules with flexible conformational states, where a fixed FRET efficiency or dye-dye distance does not provide a reasonable approximation. The comparison between the Langevin and non-Langevin models here was not to show the superiority of the Langevin method over the non-Langevin method as the Langevin method is considered an improvement simply because it is more realistic. Instead, the comparison was made to show the newly added Langevin model can be recovered from the data using typical data analysis methods at least as accurately as the original non-Langevin model and is this compatible with the PyBroMo software.

In the results presented above, the data from two example simulations using the Langevin and the non-Langevin methods were analyzed using some typical methods applied to experimental smFRET data. In the first example, a simple model for a flexible molecule where the dye-dye distances evolve dynamically using a Langevin simulation method with a harmonic free energy, generated a distribution of distances and FRET efficiencies in a physically justifiable way. In the second example, the same Langevin simulation method generated dye-dye distances with bistable free energy to model a system that inter-coverts between two states. Both examples are compared with simulated data generated with non-Langevin methods for single and bistable states with other parameters that match the Langevin simulations as closely as possible. This is done as a validation exercise to identify any unintended artifacts from the new new conformational heterogeneity when compared with the non-Langevin data using standard analysis methods including applying photon count thresholds, binning data over 1 ms, creating histograms, and fitting HMMs to estimate state transitions. Also, this allows for a comparison of some commonly used freely diffusing smFRET data analysis techniques to detect differences in the internal conformational dynamics.

Our results demonstrate both, agreement between the Langevin and non-Langevin results as well as reasonable accuracy in reproducing some of the major parameters of the underlying simulation. For instance, the histogram analyses reproduced the locations of efficiency peaks used as Langevin simulation parameters, in approximately equal proportions for the dye-dye distance distributions. Additionally, the HMM estimated similar transition matrices for the Langevin and non-Langevin timestamp data. Both Langevin and non-Langevin data agree quantitatively with the ground truth within the predicted error limits for their respective analyses.

Qualitative differences are present between the non-Langevin and Langevin timestamp data in the histogram analysis. The histograms of the Langevin timestamp data showed broader distributions of the efficiency states, in general. The comparatively narrow distributions of efficiencies from the non-Langevin timestamp data were due solely to the Brownian motion of the molecule through the PSF, but the underlying efficiency distributions are point masses. Both Langevin and non-Langevin simulation methods contained the same Brownian motion and PSF parameters so any broadening of the efficiency distribution for the Langevin timestamp data can be attributed to the ensemble of dye-dye distances from the Langevin dynamics. Similarly, the BVA also showed differences in both the positioning of the two major peaks and the population of the inter-converting states. According to the BVA analysis, we believe that Langevin data has more inter-conversions between states and also its two major states are slightly dynamic due to local protein conformational fluctuations.

It is of note that the conversion between efficiency and distance, as done in the histogram analysis, is potentially impacted by averaging of time bins and generally non-linear. We see from the agreement in Figure 3 that the averaging over 1 ms of simulated data gives a reasonable approximation of the underlying dynamics. Qualitative observations, like relative peak heights, can change after conversion. This is most obvious in Figure 4, where the two FRET states have different efficiency peak heights but the peaks of distance histograms (and underlying distribution for the Langevin simulation) are the same height. The two efficiency models used in this paper have qualitative similarities but each model required its own conversion. FRET is most accurate near the *R*_0_ value for the dye pair, with efficiency data becoming more distorted as it approaches zero or one. Accurate conversion of efficiency histogram states into distance is required to infer the underlying state information.

Beyond validation, the qualitative similarity in results from analysis of both non-Langevin and Langevin data implies the need for more sophisticated analysis methods. Despite the stark differences in the ground truth of dye-dye distances, it would be difficult to identify the Langevin results from the non-Langevin results when presented in isolation. Some identifiers of the underlying ground truth are present, like the wider spread of apparent efficiencies, but that is only visible with a direct comparison and could be missed if viewed alone. Burst analysis techniques like BVA and FRET-2CDE provide some of the advanced tools needed to distinguish static states from interconverting states. Again, the small differences are visible in a direct comparison but they are subtle and might not be apparent in isolation.

The conventional histogram analysis methods we applied to the timestamp data used time bins to collect the individual detected photons into an aggregate signal. An aggregate signal is necessary to collect enough FRET signal to overcome the background noise. For the Langevin simulation method, the time bins contain photons with an underlying ensemble of dye-dye distances and efficiencies, but the ensemble becomes averaged over the time of each bin. This is especially true when the underlying dynamics are significantly faster than the bin size. Reducing the size of time bins may reduce the averaging of conformations but also increases the proportion of background noise relative to the smFRET signal. A balance between time bin length and background noise limits how short the time bins can be while containing significant photon counts.

Using Langevin dynamics to include conformational heterogeniety, researchers will have the ability to repeatedly generate large amounts of data with a known ground truth of heterogeneous dye-dye distances. Different simulation parameters can easily be changed to generate timestamps and test assumptions based on experimental diffusing smFRET data of flexible molecules with heterogeneous states. New analysis methods beyond the standard time bin methods can then be developed and tested against the simulated data with a known ground truth to assess the effectiveness of such approaches with the ultimate goal of extracting more information from diffusing smFRET experiments of flexible molecules.

## Conclusion

In conclusion, we have shown that the addition of Langevin dynamics to freely-diffusing smFRET simulations is capable of generating timestamp data with more realistic heterogeneity of dye-dye distance dynamics and distribution. The purpose of this manuscript was to show how protein conformational dynamics can be added to a typical smFRET data simulator. We compared the Langevin and non-Langevin models to show that in both cases, it is possible to use common sm-FRET data analysis techniques to produce data consistent with their corresponding ground truth to a reasonable extent. The non-Langevin model is useful as long as the protein conformational dynamics occurs on a sub-millisecond timescale. Otherwise, Langevin model will be more practical. It is important to note that there are various other aspects of smFRET experiments that are not considered here. For instance, an explicit model to consider dye rotational/linker dynamics can be added to the proposed simulator. The implementation of the Langevin dynamics provides a flexible approach for defining the underlying dynamics of the molecule with full knowledge of the ground truth. Simulated data with known ground truth of realistic heterogeneous dye-dye distances will play an important role in developing new techniques for the analysis of freely diffusing smFRET data for flexible molecules.

## Author Contributions

Authors J.L., A.P., M.J., A.C.C., and H.W. performed the research and analysis. Authors J.L., S.S., A.P., M.J., A.C.C., H.W., D.S.M, R.R., and M.M wrote the manuscript. Authors R.R., D.S.M, and M.M. designed the research.

## Data Availability

Simulation and analysis scripts are available on GitHub Page: https://github.com/bslgroup/ PyBroMo

## Acknowledgement

This research is supported by the National Science Foundation under Awards 1940188, 1945465, 1934985, 1940124, and 1940179. This research is also supported by the Arkansas High Performance Computing Center which is funded through multiple National Science Foundation grants and the Arkansas Economic Development Commission. shown in this document.

## Supporting Information Available

Includes additional details of Simulation software and Example experimental smFRET analysis including quantifying error for HMM analysis, changepoint analysis, BVA: single state analysis and FRET-2CDE analysis.

## TOC Graphic

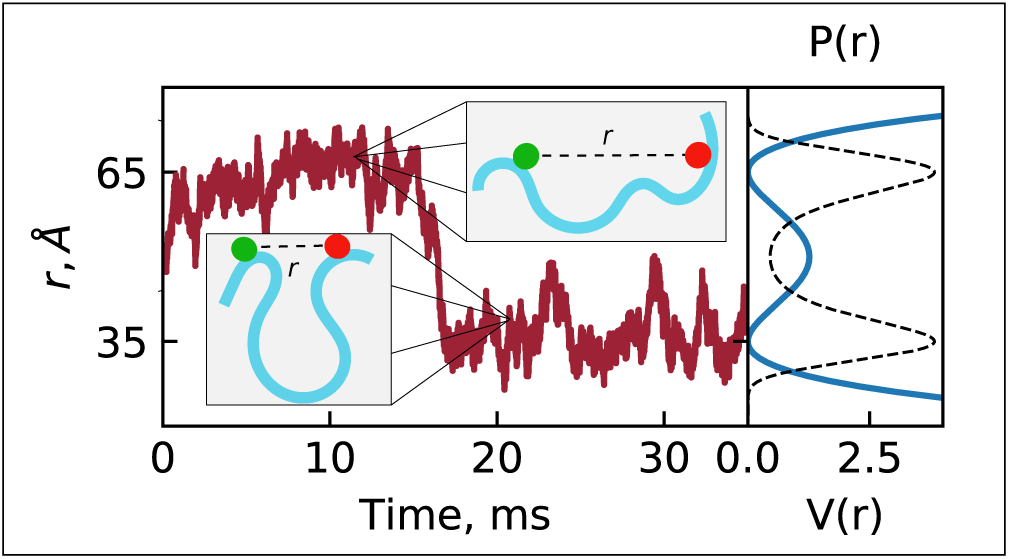

