## Supporting Information for "Simulating Freely-diffusing Single-molecule FRET Data with Consideration of Protein Conformational Dynamics"

#### Simulation software

PyBroMo[1] simulates photon emission from fluorescent dye pairs attached to freely-diffusing molecules while recording the timestamps from those emissions, similar to experimental FRET data. This software was designed to generate realistic FRET data by handling multiple populations of molecules with their own diffusion coefficients and FRET efficiencies, as well as generating background photons with separate emission rates for the donor and acceptor channels. This software uses a Brownian motion simulation to model the molecular diffusion in a solution, a numerical point spread function (PSF) to model the laser, and Poisson background noise to model background photon rates for each channel.

The first step in generating FRET timestamps is defining the basic elements of the simulation in the form of a Python script. In the script, the molecules are defined by a population number and diffusion coefficient, $D_B$. The simulation is defined by providing the number of molecules and the simulation box dimensions, $L_x, L_y$, and $L_z$, as well as conditions for how to handle molecule interactions with the boundary. If a molecule's position is advanced across the box boundary, the position is either wrapped across the opposite boundary, or reflected back across the same boundary that was crossed. A point spread function (PSF) is defined to model the laser focal beam inside the simulation box. The PSF represents the emission probability of a molecule at any position within the simulation box. A Gaussian PSF is available where the emission probability in all dimensions is defined by

$$f(x) = \frac{1}{\sigma_x \sqrt{2\pi}} e^{-\frac{1}{2}\left(\frac{x-\mu_x}{\sigma_x}\right)^2} \tag{1}$$

where $\mu_x$ is the mean coordinate for the center of the function and $\sigma_x$ is the standard deviation. Eq. (1) can be extended to include other Cartesian coordinates $y$ and $z$. PyBroMo is also capable of importing custom PSF functions from tools like PSFLab[2] that can generate a custom numerical PSF that includes factors like light polarization. PyBroMo includes a default numeric PSF for use without the user having to create their own.

Next, the simulation inputs are passed to the Brownian motion simulation module along with a timestep ($\delta t$) and a maximum time to advance the molecules through the simulation box. The Brownian motion is a stochastic process where the position in each dimension are advanced from the current position by a random number drawn from a normal distribution,

$$x(t + \delta t) = \quad x(t) + \xi \tag{2}$$

where $\xi \sim N(0, 2D_B\delta t)$ for the white noise contribution. The Brownian motion simulation then repeatedly advances each molecule's position in three dimensions by the $\delta t$ until the maximum time is reached. At each time step, the PSF calculates the normalized emission probability for every molecules position in a trajectory vector, $\mathscr{P}$. Molecules in regions of high emission probability, near the center of the PSF, emit more photons up to the maximum emission rate.

Finally, the timestamp generation module creates the number of photon emissions events, $\kappa$, through a discrete random Poisson process

$$f(\kappa,\lambda) = \frac{\lambda^{\kappa} e^{-\lambda}}{\kappa!} \tag{3}$$

where $\lambda$ is the expected number of emissions. The values needed to calculate the $\lambda$ values for every time step are a maximum total emission rate, $\varepsilon_T$, efficiency, $E$, for each population, and the emission probabilities, $\mathscr{P}$ from the Brownian motion simulation. Emission rates for the acceptor, $\varepsilon_{Acc}$, and donor , $\varepsilon_{Don}$, channels are then calculated

$$\varepsilon_{Acc} = \varepsilon_T E \tag{4}$$

$$\varepsilon_{Don} = \varepsilon_T (1-E) \tag{5}$$

The efficiency, $E$, is constant for all timesteps. Separate expected counts for the acceptor, $\lambda_{Acc}$ and donor, $\lambda_{Don}$ are then calculated

$$\lambda_{Acc} = \mathscr{P}\varepsilon_{Acc}\delta t \tag{6}$$

$$\lambda_{Don} = \mathscr{P}\varepsilon_{Don}\delta t \tag{7}$$

and used to randomly draw emission events at every time step. Similarly, background emissions rates are also determined for the acceptor and donor detector channels by randomly drawn numbers from a Poisson distribution with expected values, $\lambda_{BGAcc}, \lambda_{BGDon}$, supplied as a simulation parameter.

The timestamps are merged and sorted into a single trajectory for output. A vector of labels is also generated to label the timestamp as being from the acceptor or donor channel. Other values of interest that may be included are the molecule ID that generated the photon emission or the position of the molecule in the PSF.

### Example Experimental smFRET Analysis

As an example of a standard analysis used with experimental data, we include a histogram and Gaussian mixture analysis of real freely diffusion smFRET data for the C-terminus of the Albino3 protein (CAlb3)[3], an intrinsically disordered protein.

The results appear in Figure 1. Plot A shows a trace of the combined acceptor and donor

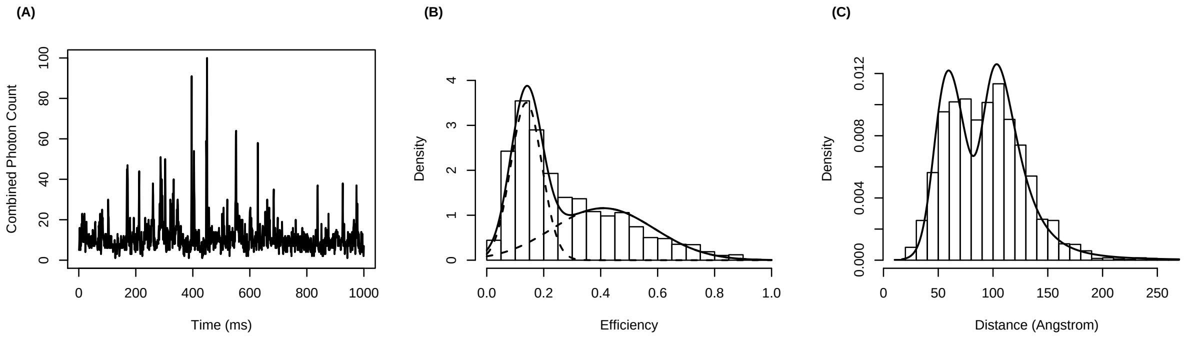


Figure 1: (A) Trace plot of the combined acceptor and donor counts from a one second segment of the cAlb experiment. (B) The estimated Gaussian mixture density (solid black line) on top of a histogram of the apparent efficiencies. (C) The corresponding plot in the distance space.

counts from a one second segment of the 10 second experiment. Plot B shows the estimated two-component Gaussian mixture density on top of a histogram of the apparent efficiencies. The low efficiency component has a mean of 0.14, a standard deviation of 0.05, and a mixture weight of 0.48, while the high efficiency component has a mean of 0.41, a standard deviation of 0.18, and a mixture weight of 0.52. Plot C shows the corresponding histogram and density after a transformation to the distance space.

#### Quantifying error for HMM analysis

The true transition matrix can be computed theoretically as,

$$T^{(0)} = \begin{pmatrix} 0.968 & 0.032 \\ 0.032 & 0.968 \end{pmatrix}. \tag{8}$$

The error between the measured and theoretical transition matrices can be quantified in terms of a weighted Kullback-Leibler divergence,[4,5] as

$$D(T^{(0)}||T^{\mathrm{L}}) = \sum_i \pi_i(T^{(0)}) \sum_j T_{ij}^{(0)} \log \left( \frac{T_{ij}^{(0)}}{T_{ij}^{\mathrm{L}}} \right) = 0.0036, \tag{9}$$

where $\pi(T)$ is the equilibrium distribution vector based on the transition matrix $T$, which satisfies $\pi T = \pi$, which for the symmetric matrix $T^{\mathrm{L}(0)}$ is $\pi(T^{\mathrm{L}(0)}) = (0.5, 0.5)$

We split our binned photon count data in 10 parts of equal length. We perform the same procedure, obtaining a set of bursts through thresholding, and fitting an HMM to each set of bursts, for the 10 data subsets. This way we extract 10 measured transition matrices, $T^i$, with $i = 1, \ldots, 10$. With this set of transition matrices, we define the average transition matrix as

$$\bar{T} = \begin{pmatrix} \frac{\sum_{i=0}^{10} T_{11}^i}{(\sum_{i=0}^{10} T_{11}^i + \sum_{i=0}^{10} T_{12}^i)} & \frac{\sum_{i=0}^{10} T_{12}^i}{(\sum_{i=0}^{10} T_{11}^i + \sum_{i=0}^{10} T_{12}^i)} \\ \frac{\sum_{i=0}^{10} T_{21}^i}{(\sum_{i=0}^{10} T_{21}^i + \sum_{i=0}^{10} T_{22}^i)} & \frac{\sum_{i=0}^{10} T_{22}^i}{(\sum_{i=0}^{10} T_{21}^i + \sum_{i=0}^{10} T_{22}^i)} \end{pmatrix}, \tag{10}$$

where the normalization factors for each coefficient are included to ensure each row of the transition matrix is a proper probability distribution.

We can now compute the error between each of the transition matrices, $T_i$, and the average matrix, $\bar{T}$, which gives us information on the precision of our procedure, in particular we characterize the precision by computing the average deviation,

$$\sigma_{\bar{T}}^{\mathrm{L}} \equiv \frac{1}{10} \sum_{i=1}^{10} D(\bar{T}||T_i) = 0.00015. \tag{11}$$

We similarly compute the average deviation between the known transition matrix and each of the ten transition matrices

$$\sigma_T^{L,(0)} \equiv \frac{1}{10}\sum_{i=1}^{10} D(T^{(0)}||T_i) = 0.0037. \tag{12}$$

Comparing $\sigma_{\bar{T}}$ and $\sigma_{T^{L(0)}}$ shows that the error between our measured and true transition matrices is not completely explained by the variability of the measured transition matrices, thus the error is unlikely to be reduced by simply analyzing more data.

We perform the same analysis on the Non-Langevin data, for comparison. Here the error between the measured and true transition matrices is given by,

$$D(T^{(0)}||T^{\mathrm{NL}}) = 0.0053\,. \tag{13}$$

We again quantify the variability of the measured transition matrices by splitting the data in 10 parts, and measuring a transition matrix for each. The average deviation between each of these matrices and the average of the ten transition matrices is

$$\sigma_{\bar{T}}^{\mathrm{NL}} = 0.00020, \tag{14}$$

and the average deviation between each of the ten matrices and the known transition matrix is,

$$\sigma_T^{NL,(0)} = 0.0056\,. \tag{15}$$

While showing qualitatively similar behavior, our analysis shows smaller measures of error for the Langevin data, compared to the non-Langevin case.

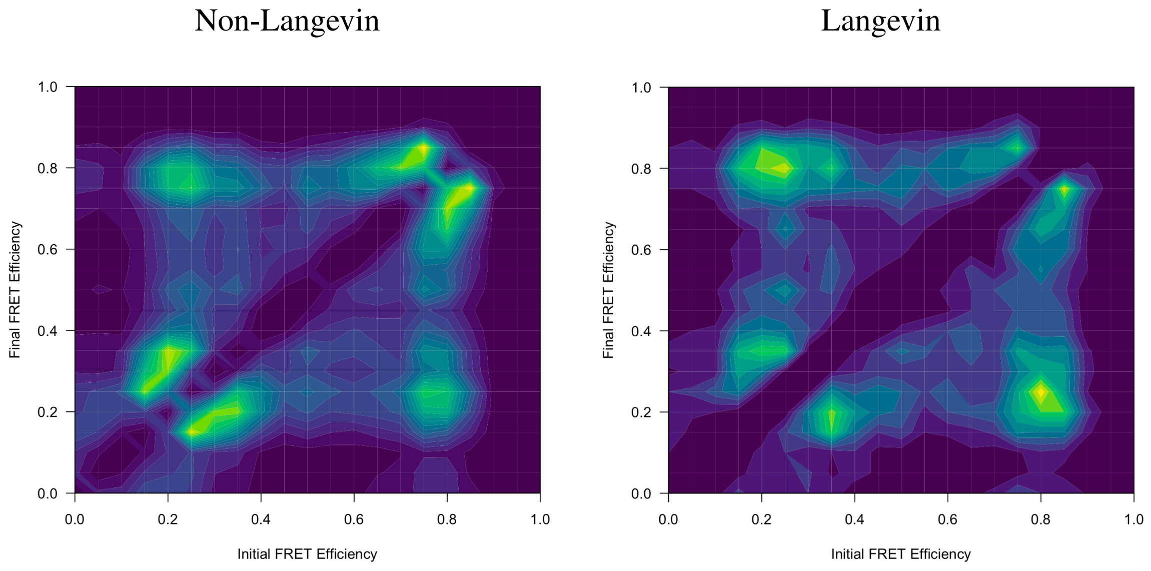


Figure 2: Heatmap of count of apparent efficiency state changes within both Langevin and Non-Langevin simulated data from ABCO changepoint algorithm. The two peaks away from the diagonal indicate inter-state changes between the two states while peaks along diagonals indicate intra-state changes.

### Changepoint Analysis

Using a changepoint algorithm, we analyze the state switching dynamics of the Langevin and Non-Langevin binned data, converted into apparent efficiencies. The resulting heat map, shown in Figure 2, illustrates the state transitions densities derived from the changepoint algorithm, with initial apparent efficiency on the x-axis and final FRET apparent efficiency on the y-axis. A brighter color in the heat map indicates a higher density for transition between the two efficiencies. As seen in the figure, the two peaks describe transitions between the low smFRET state and high smFRET state in our simulated data. We see that most changes in the heatmap are reflected in these two regions. In addition, we also see small peaks along the diagonals for intra-state changes. These small peaks reflect the intra-state dynamic of the data and the variability within the low-FRET and high-FRET states. Overall, the heatmaps generated from the non-Langevin and Langevin timestamp data identify the two distinct efficiency states near the ground truth of 0.290 and 0.944. A noteworthy difference is the smoother transitions between states in the non-Langevin heat map compared with the Langevin heatmap.

To give some detail about the changepoint algorithm, we utilize Adaptive Bayesian Changepoint Analysis & Local Outlier Scoring (ABCO)[6] on the binned efficiencies. ABCO decomposes the apparent efficiencies, $\hat{E}_t$, into three components: a local trend term, $\beta_t$, a sparse, additive outlier term, $\zeta_t$, and a heteroskedastic noise process, $\varepsilon_t$,

$$\hat{E}_t = \beta_t + \zeta_t + \varepsilon_t. \tag{16}$$

The flexible outlier term, $\zeta_t$, and the error term, $\varepsilon_t$, make the model robust to noise and adapt well across time-varying dynamics. A dynamic horseshoe prior[7] is placed on the first difference of $\beta_t$ to ensure that most changes in $\beta_t$ will be small while allowing for large jumps in certain timestamps. These large jumps timestamps, calculated as changes exceeding a estimated threshold, will be classified as changepoints. For each changepoint, the initial and final Fret efficiencies are stored and used to generate the heat map shown in Figure 2.

### BVA: Single State

A BVA analysis on the data from Example 1 is shown in Figure 3. The shorter trajectory length made for fewer bursts and robust statistical information, when using the same definition of a burst as found in the main paper. Nevertheless, both the non-Langevin and Langevin data show similar plots with relatively little variance between the peaks.

### FRET-2CDE

Another burst metric called FRET-2CDE measures the density of photons surrounding each photon in a burst to quantify variations in emission rates. The density of a photon is considered respective to its own channel and the opposing channel in the case of two color FRET. The FRET-2CDE metric is designed to be centered around 10 with higher scores indicating potential areas of heterogeneity in photon distributions. The values for FRET-CDE for the non-Langevin and Langevin

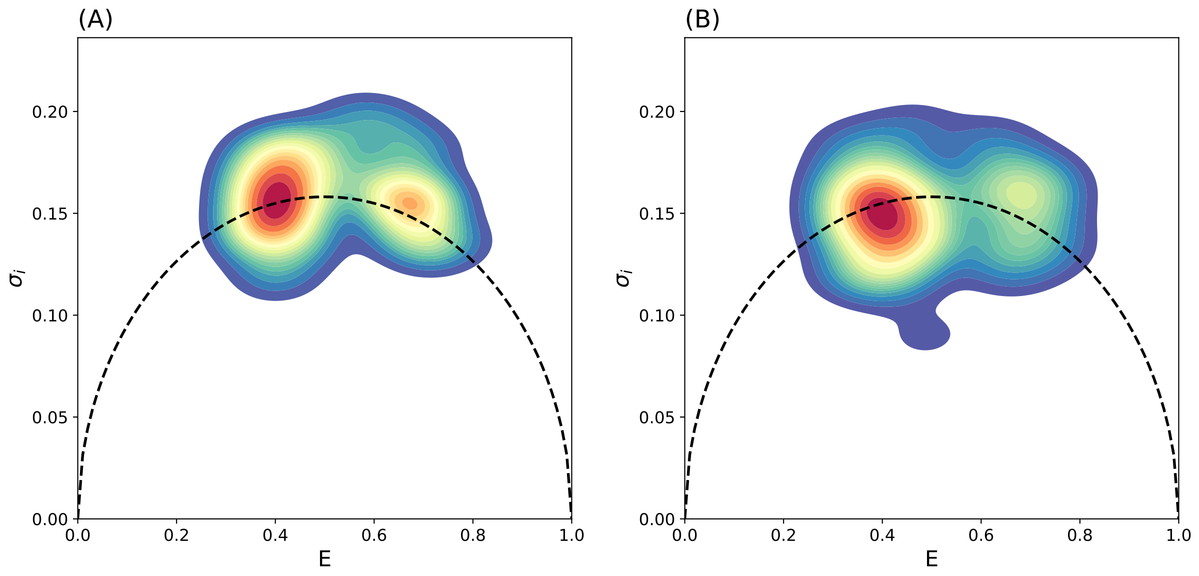


Figure 3: BVA using the Example 1 data to compare the (A) non-Langevin and (B) Langevin data. A KDE is used to present smooth contours the scatter plot of efficiency and variance of efficiency for each burst. The dotted line shows the standard deviation of the binomial distribution for 10 samples.

data sets are show in Figure 5, again as a contour plot from a KDE of the data.

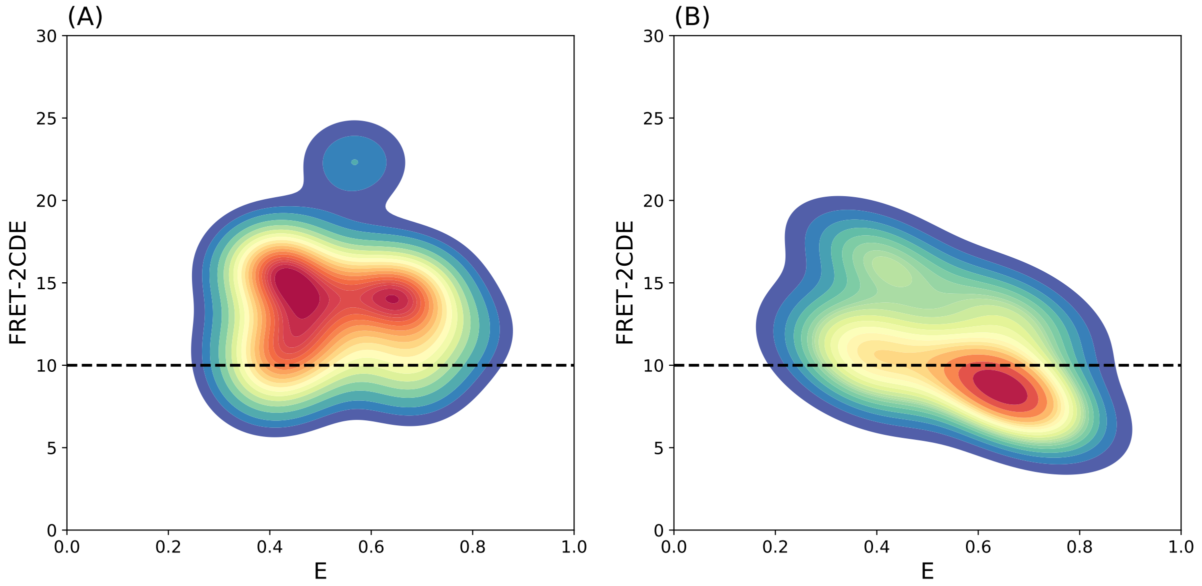


Figure 4: The efficiency and FRET-2CDE values represented as a scatter plot using a KDE to compare the (A) non-Langevin and (B) Langevin data from the single state data. The dotted line on 10 represents the FRET-2CDE value when the neighboring photons are distributed around a referent photon.

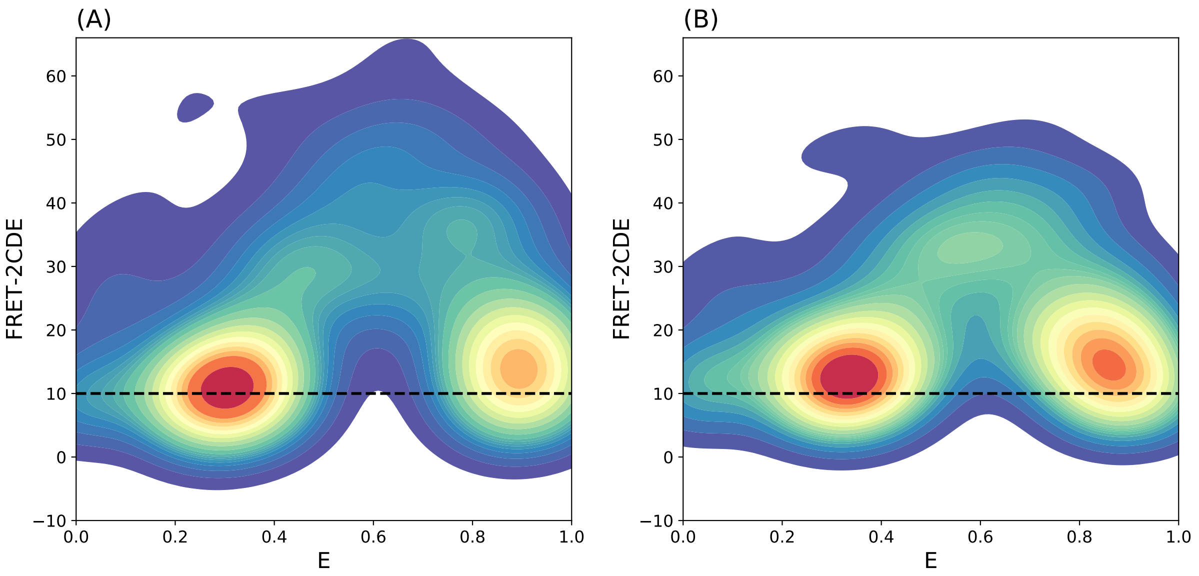


Figure 5: The efficiency and FRET-2CDE values represented as a scatter plot using a KDE to compare the (A) non-Langevin and (B) Langevin from two state inter-conversion data. The dotted line on 10 represents the FRET-2CDE value when the neighboring photons are distributed around a referent photon.
